# Strain-dependent impact of G and SH deletions provide new insights for live-attenuated HMPV vaccine development

**DOI:** 10.1101/781302

**Authors:** Julia Dubois, Andrés Pizzorno, Marie-Hélène Cavanagh, Blandine Padey, Claire Nicolas de Lamballerie, Olus Uyar, Marie-Christine Venable, Julie Carbonneau, Aurélien Traversier, Thomas Julien, Christian Couture, Bruno Lina, Marie-Ève Hamelin, Olivier Terrier, Manuel Rosa-Calatrava, Guy Boivin

## Abstract

Human metapneumovirus (HMPV) is a major pediatric respiratory pathogen with currently no specific treatment or licensed vaccine. Different strategies to prevent this infection have been evaluated, including live-attenuated vaccines (LAV) based on SH and/or G protein deletions. This approach showed promising outcomes but has not been evaluated further using different viral strains. In that regard, we previously showed that different HMPV strains harbor distinct *in vitro* fusogenic and *in vivo* pathogenic phenotypes, possibly influencing the selection of vaccine strains. In this study, we investigated the putative contribution of the low conserved SH or G accessory proteins in such strain-dependent phenotypes and generated recombinant wild type (WT) and SH- or G-deleted viruses derived from two different patient-derived HMPV strains, A1/C-85473 and B2/CAN98-75.

The ΔSH and ΔG deletions led to different strain-specific phenotypes in both LLC-MK2 cell and reconstituted human airway epithelium models. More interestingly, the ΔG-85473 and especially ΔSH-C-85473 recombinant viruses conferred significant protection against HMPV challenge and induced immunogenicity against a heterologous strain. In conclusion, our results show that the viral genetic backbone should be considered in the design of live-attenuated HMPV vaccines, and that a SH-deleted virus based on the A1/C-85473 HMPV strain could be a promising LAV candidate as it is both attenuated and protective in mice while being efficiently produced in a cell-based system.

## 1. Introduction

Human metapneumovirus (HMPV) is a worldwide cause of acute respiratory tract infections (ARTI) among children, the elderly and immunocompromised individuals [1, 2]. HMPV infections share many features with those of the human respiratory syncytial virus (HRSV), also belonging to the *Pneumoviridae* family [3, 4]. Despite the important clinical burden in infants and young children, no licensed vaccine or specific and potent antiviral are currently available. While several HRSV vaccine candidates have already entered clinical trials [5], some HMPV candidates have shown the potential to progress towards clinical evaluation stages [6]. In that regard, different HMPV vaccine strategies have been evaluated in animals, from formalin-inactivated vaccine, leading to enhanced disease [7], to the elaboration of protein-based recombinant vaccines or live-attenuated vaccines (LAV) [6]. Among them, LAV have shown the potential to elicit both humoral and mucosal immunity and mimic natural viral replication routes, and they are therefore considered as highly suitable for HMPV pediatric immunization strategies [8].

HMPV viruses are divided into two main phylogenetic lineages (A and B), which are further divided into at least two sub-lineages (A1, A2a/A2b, B1 and B2) [9-13]. The HMPV genome is composed of a negative single stranded RNA molecule of approximately 13 kb in length, containing eight genes encoding for nine different proteins [14, 15], including three surface glycoproteins (F, G, SH). The F glycoprotein is the major HMPV antigen [16] and leads to both attachment and fusion of viral particles to the target cell [17]. In contrast, the exact role of G and SH glycoproteins is still a matter of debate. Indeed, the F protein of HMPV has been shown to bind not only the cellular integrin αVβ1 receptor but also glycosaminoglycans (GAGs), such as heparan sulfate, hence being able to substitute to the virus GAG-mediated attachment function once exclusively attributed to the G protein [18-21]. Nonetheless, a role of the G protein was also suggested in the host cell response to infection [22-24]. For instance, stimulation of the retinoid-acid inducible gene 1 (RIG-I) signaling pathway was reported with a recombinant HMPV (rHMPV) lacking the G protein (ΔG) *in vitro*, which led to increased NF-κB activation and enhanced cytokine secretion [22]. On the other hand, the SH protein has been shown to alter the NF-κB pathway [25] and also to form a viroporin complex at the cell membrane [26].

The G and SH proteins have been considered for a long time as “accessory” non-essential proteins for HMPV replication [27], as illustrated by recombinant HMPV viruses lacking either G, SH or both genes that can replicate efficiently *in vitro* and *in vivo* [28]. Moreover, the contribution of G and SH proteins to HMPV antigenicity, as well as the attenuation phenotype associated to G-deleted virus, led to consider such modified viruses as potential LAV candidates [16, 27]. However, the achievement of such perspectives is nuanced by the fact that all previous studies were based on a unique HMPV backbone, notably the prototypical CAN97-83 strain from the A2 sub-lineage. In that regard, many HMPV subtypes co-circulate each year with high genetic diversity among A and B subtypes, particularly in the case of the less conserved G and SH proteins, with approximately 37% and 59% amino acid sequence homologies between subtypes, respectively [9, 14]. In parallel, we and others previously demonstrated that HMPV viruses diverge in their *in vitro* and/or *in vivo* phenotypes in a strain-dependent manner, notably by considering the HMPV F and G proteins and their functions [18, 29-33].

In this context, we generated recombinant wild type (WT) and SH- or G-deleted viruses (ΔSH and ΔG respectively) from two patient-derived HMPV strains (A1/C-85473 and B2/CAN98-75) and compared the respective functional impacts of SH and G deletions. In this context, we observed different strain-specific phenotypes both in LLC-MK2 cells and reconstituted human airway epithelium (HAE) models that provided new insights on the importance of the genetic background in the design of HMPV LAV. This prompted us to evaluate the ΔSH-C-85473 and ΔG-C-85473 recombinant viruses in BALB/c mice for their capacity to induce efficiently neutralizing antibody response, recruit immune cells in lungs and protect mice against lethal HMPV challenge. Our results suggest that an HMPV C-85473 strain-based LAV harboring SH and/or G deletions would be a promising platform for further development of pneumovirus vaccine candidates.

## 2. Materials and Methods

### Cells

LLC-MK2 cells (ATCC CCL-7) were maintained in minimal essential medium (MEM, Life Technologies) supplemented with 10% fetal bovine serum (FBS, Wisent) and 10 mM HEPES (Sigma). BSR-T7/5 cells (a kind gift from Dr Ursula Buchholz at the NIAID in Bethesda, MD) were cultured in MEM supplemented with 10% FBS, 1% Non-essential amino acids (NEAA) (Life Technologies), 10 mM HEPES (Sigma), 1% penicillin/streptomycin (Wisent) and 0.2 mg/ml geneticin (G418, Life Technologies).

### Rescue of recombinant viruses

pSP72 plasmids (Promega) encoding the full-length genomic cDNA of HMPV B2/CAN98-75 or A1/C-85473 strains (GenBank accession numbers: AY145289.1 and KM408076.1, respectively) were constructed as previously described [29, 34]. To generate G- or SH-deleted viruses, plasmids encoding HMPV antigenomes were amplified with Phusion DNA polymerase (New England Biolabs) using primers (listed in **Table 1)** designed to match with the 3’ and 5’ regions of either G or SH genes, and taking care to conserve an intergenic sequence between flanking genes (as represented in **Figure 1A**). After amplification, linear DNA was phosphorylated and re-ligated by T4 Ligase. All plasmids were sequenced completely.

**Table 1.**
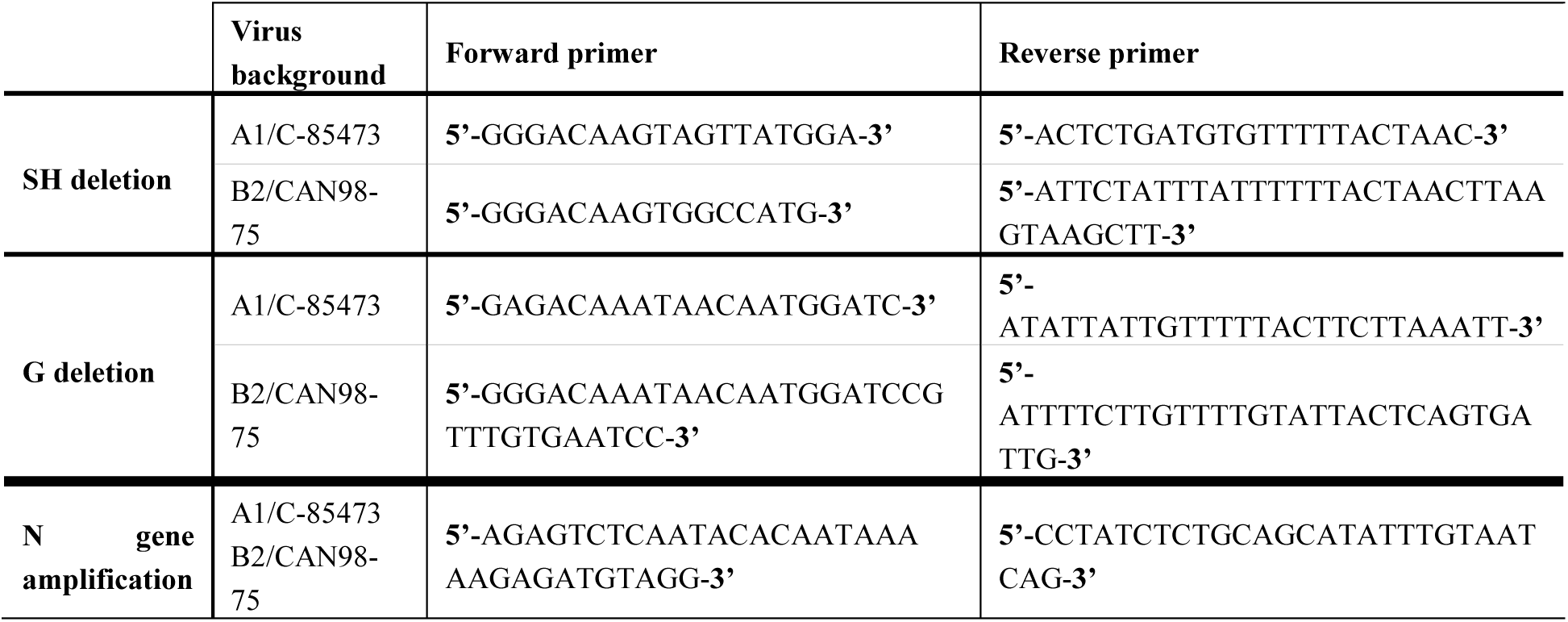
List of primers

**Figure 1:**
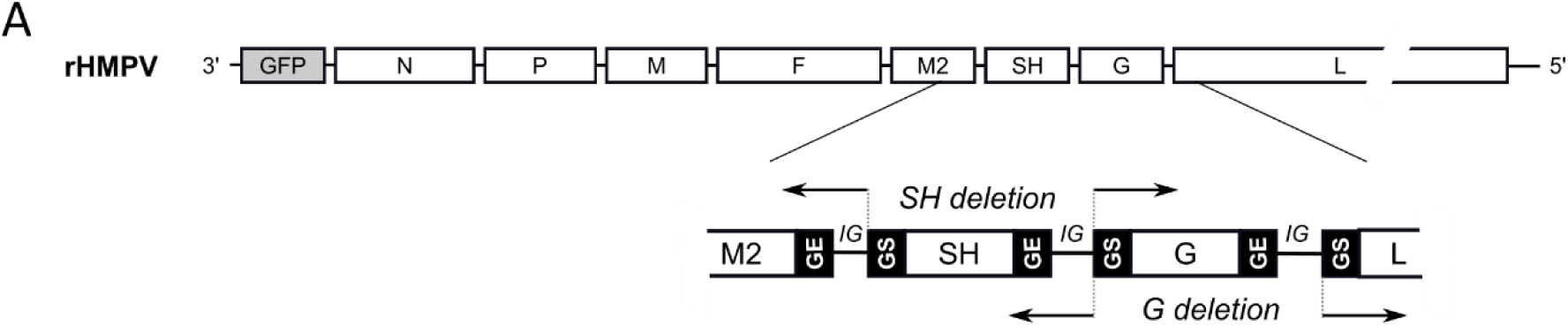
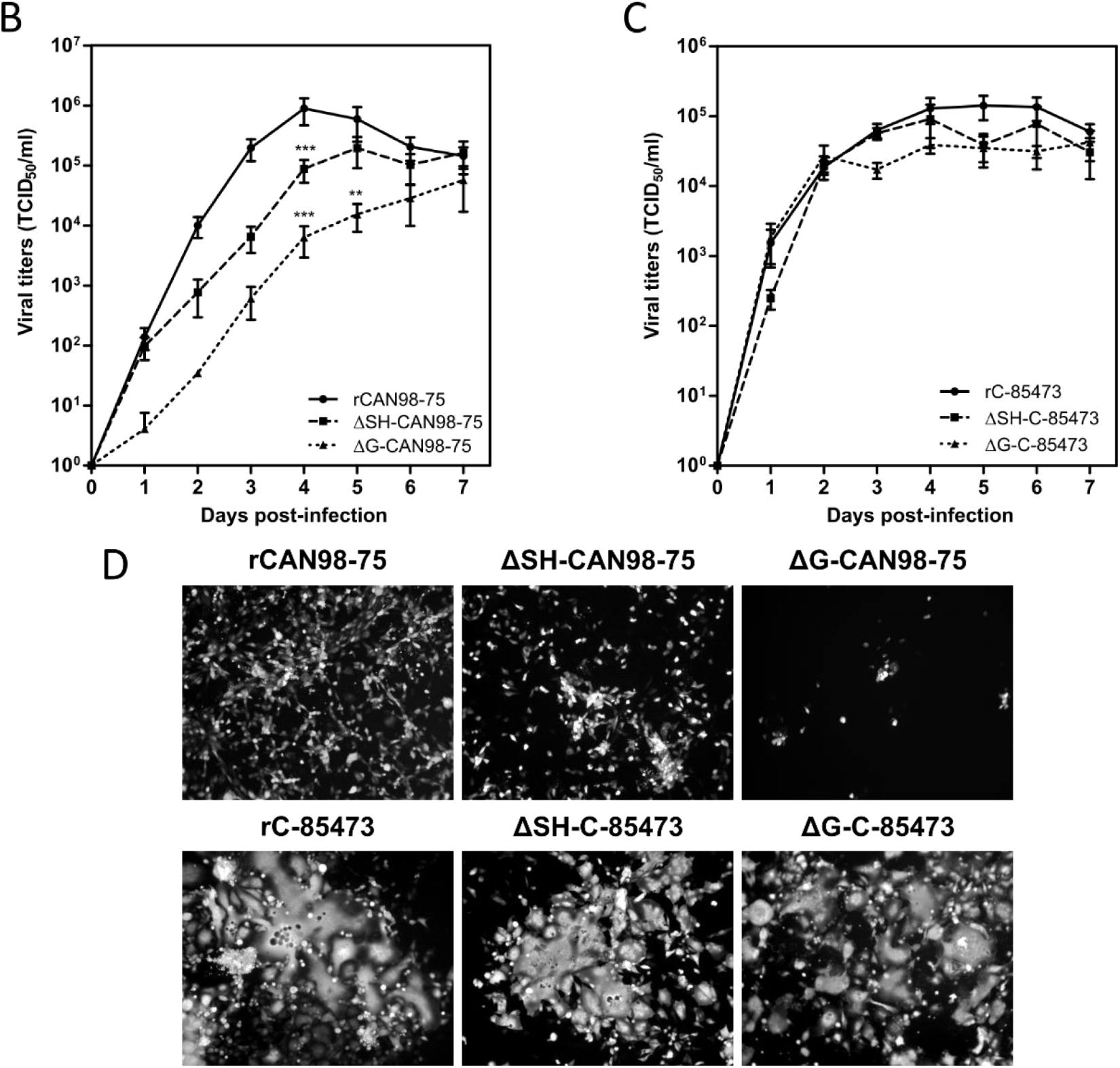
Construction of recombinant HMPV viruses and *in vitro* replicative capacity of SH and G-deleted rHMPV viruses. **(A)** Schematic representation of the genome of a recombinant GFP-expressing rHMPV viruses, with specific focus on the region of the genome describing the position of primers used for the deletion strategy (arrow, table 1). GS: Gene Start. GE: Gene End. IG: intergenic sequence. **(B-C)** LLC-MK2 monolayers in 24 wells-plates were infected with each of the 6 recovered rHMPV strains (rCAN98-75, ΔSH-CAN98-75, ΔG-CAN98-75; rC-85473, ΔSH-C-85473, ΔG-C-85473) at a MOI of 0.01. Supernatants were harvested every 24 h for 7 days, frozen, sonicated and titrated as TCID_50_ / ml on LLC-MK2 cells. Growth curves represent mean titers of 2 independent experiments, with each time point titrated in triplicate. ** p < 0.01, *** p < 0.001 when comparing each ΔSH / ΔG virus to its WT counterpart using Repeated Measures Two-way ANOVA. **(D**) Images of representative cytopathic effects and spread of each virus at 3 days post-infection were captured using fluorescent microscopy (10x magnification).

BSR-T7 cells were transfected with the HMPV antigenome constructions plus 4 supporting plasmids, encoding N, P, L, and M2-1 ORFs of CAN98-75 and LLC-MK2 cells were added for co-culture (2 to 3 days), as previously described [29]. Cells were harvested, sonicated, centrifuged and the supernatant was diluted to inoculate LLC-MK2 monolayers. After 3 to 4 amplification passages onto LLC-MK2 cells, recombinant GFP-expressing viruses (rHMPV) were concentrated by ultracentrifugation, resuspended in OptiMEM (Life Technologies) and stored at −80°C. Viral stocks were titrated as 50% tissue culture infectious doses (TCID_50_)/ml [35].

### *In vitro* experiments

For replication kinetics assays, confluent monolayers of LLC-MK2 cells in 24-wells plates were infected with rHMPV viruses at a MOI of 0.01, as described previously [29]. Supernatants of infected wells were harvested in triplicate every 24 h for 7 days and end-point TCID_50_/ml titrations were performed on each sample.

For viral binding assays, confluent LLC-MK2 monolayers were washed with cold PBS and placed on ice before rHMPV inoculation at MOI 0.5. After adsorption times on ice (30 min, 1h, 2h or 3h), the inoculum was removed and replaced by fresh infection medium. After a 24h incubation at 37°C and 5% CO^2^, cells were trypsinized, centrifuged and fixed using 2% formaldehyde. The percentages of infected GFP positive cells over 1×10^4^ total cells were measured by flow cytometry by using a FACSCantoII analyzer (Becton Dickinson) with FACSDiva software. Data were analyzed by using the FlowJo software (Tree Star, Inc.) and normalized to each virus inoculum.

For experimental virus entry assays, confluent LLC-MK2 cells were inoculated at a MOI of 0.5 after being placed on ice for 5 min and washed with cold PBS. The binding of virus to cells proceeded for 2 h on ice until the inoculum was replaced by fresh infection medium. Cells were incubated for 5 min, 30 min, 1 h or 2 h at 37°C and 5% CO2 to allow virus entry into cells, medium was then removed and citrate buffer (40 mM sodium citrate dihydrate, 10 mM potassium chloride, 135 mM sodium chloride, pH 3.0) was added to inactivate any remaining extracellular virus, as previously described [38]. Cells were washed in PBS and incubated at 37°C in fresh medium during 24 h. Infected cells were quantified by GFP detection in flow cytometry and values were normalized to condition without viral inactivation.

### Infection of reconstituted human airway epithelium

*In vitro* 3D reconstituted human airway epithelium (HAE), derived from a pool of healthy donors’ primary bronchial cells (MucilAir™) were purchased from Epithelix (Switzerland). A viral inoculum corresponding to a MOI of 0.1 was added onto HAE and incubated for 1.5 h at 37°C and 5% CO2. After 5 days of infection, captured images of infected HAE by fluorescent microscopy were taken and the whole epithelium was scraped in RLT buffer (Qiagen). Total RNA was then extracted with RNeasy minikit (Qiagen) to perform viral genome quantification by RT-qPCR.

### Real-time RT-PCR

The total RNA extracted from samples was randomly reverse transcribed using SuperScriptII RT (Invitrogen) at 42°C. Amplification of the HMPV N gene was performed by quantitative PCR using SYBR Green qPCR Master Mix (Agilent) and a set of primers listed in **Table 1**. The calibration of HMPV N copies was assessed by amplification of a plasmid kindly provided by Dr Ab Osterhaus (Erasmus Medical Center, Rotterdam).

### Animal studies

Four to six week-old BALB/c mice (Charles River Laboratories), housed in groups of five per micro-isolator cage, were infected intra-nasally with 5×10^5^ TCID_50_ of rHMPV C-85473 viruses (WT, ΔSH or ΔG). As a control group, mice were mock-infected intra-nasally with OptiMEM. Animals were monitored on a daily basis during 14 days for weight loss, clinical signs, reduced activity or ruffled fur and were sacrificed if they reached 20% of initial weight loss. Mice were euthanized at 5 dpi using sodium pentobarbital and lungs were removed for the evaluation of viral titers or histopathological analysis, as described [29]. For virus titration, lungs were homogenized in 1 ml of PBS before TCID_50_ titration or N quantification by RT-qPCR.

For HMPV challenge experiments three weeks post-immunization with 5×10^5^ TCID_50_ of recombinant rC-85473 viruses, mice were challenged intra-nasally with 1×10^6^ TCID_50_ of WT rC-85473 HMPV virus (50% lethal dose). Mice immunized with OptiMEM were used as negative control. Viral titers and histopathological scores were evaluated at 5 days post-challenge as previously. Prior to immunization/infection and 21 days after challenge, blood samples were taken to evaluate the neutralizing antibodies against the recombinant virus (rC-85473) or WT patient-derived homologous C-85473 and heterologous CAN98-75 strains. Reciprocal neutralizing antibody titers were determined by end-point dilution assay, as described [7].

To quantify pulmonary inflammatory cytokine/chemokine levels in HMPV-immunized challenged mice, we euthanized mice and harvested lungs on days 1 and 5 post-challenge. Lungs were homogenized in D-PBS containing protease inhibitors to perform the cytokines quantification using a commercial multiplex mouse cytokine bead assay (Bio-Rad, Bio-Plex Pro Mouse Cytokine 23-plex Assay) according to the manufacturers’ instructions. Results were analyzed with the Luminex system (QIAGEN).

In order to analyze lung-infiltrating immune cells, mice were deeply anesthetized and perfused intracardially with D-PBS without Ca2+ and Mg2+ on days 1 and 5 post-challenge. Lungs were collected, digested with Liberase TL and processed to recover living leukocytes, as described [34]. Then, cells were incubated on ice for 40 min with a pool of antibodies (anti-CD45, anti-Ly6G, anti-CD11b, anti-CD170/Siglec-F, anti-Ly6C, anti-CD11c, anti-F4/80, anti-B220, anti-CD3ε, anti-CD4 and anti-CD8a, purchased from BD) to discriminate cell populations. The count numbers of total pulmonary leukocytes (CD45+), neutrophils (LyG6+), macrophages (SiglecF+, F4/80+), lymphocytes T (CD3ε+) CD4+, CD8+ and lymphocytes B (B220+) were measured by flow cytometry from whole living cells. Number of cells were determined with Precision Count Beads™ (BioLegend). Data acquisition and analyses were performed by a BD LSRII flow cytometer and the BD FACSDiva software.

Animal studies were approved by the SFR Biosciences Ethics Committee (CECCAPP C015 Rhône-Alpes, protocol ENS_2017_019) according to European ethical guidelines 2010/63/UE on animal experimentation and by the Animal Protection Committee of the Quebec University Health Centre (Protocol CPAC 2017-140-2) according to Canadian Council guidelines on Animal Care.

### Statistical analysis

Two-way ANOVA’s with Dunnett post-tests were used to compare data of ΔSH and ΔG HMPV to their corresponding WT virus. Statistical analyses were performed using GraphPad Prism7.

## 3. Results

### In vitro characteristics of ΔG- and ΔSH-HMPV recombinant viruses differ depending on the viral strain background

Based on our previous studies [29, 30], we successfully generated by reverse genetics different GFP-expressing HMPV recombinant viruses derived from the A1/C-85473 strain (rC-85473, ΔG-C-85473, ΔSH-C-85473) and the B2/CAN98-75 strain (rCAN98-75, ΔG-CAN98-75 and ΔSH-CAN98-75) (**Figure 1A**). Rescued recombinant viruses were firstly assessed for their replicative capacity in LLC-MK2 cells over a 7-day period (**Figure 1B-C**). While rCAN98-75 virus peaked at 8.95×10^5^ TCID_50_/ml at 4 dpi, ΔSH-CAN98-75 virus showed reduced replication as early as 2 dpi, with a 24 h delay to reach peak titer (1.95×10^5^ TCID_50_/ml) (**Figure 1B**). A significantly more attenuated phenotype was observed with the ΔG-CAN98-75 virus, for which the highest viral titer measured (5.72×10^4^ TCID_50_/ml at 7 dpi) was delayed and 1.2 log10 lower than that of the WT (**Figure 1B**). This differential replication among the three CAN98-75-based recombinant viruses was also illustrated by representative fluorescent microscopy images of infected LLC-MK2 cells at 3 dpi, notably considering the extent of viral spread and GFP positive cells (**Figure 1D**). On the contrary, no such important differences in replication properties were observed between the two deleted C-85473-derived recombinant viruses and their WT counterpart (**Figure 1C**). Peak viral titers of approximately 1×10^5^ TCID_50_/ml for both the C-85473-WT and ΔSH-C-85473 viruses, or a peak of 4.22×10^4^ TCID_50_/ml for the ΔG-C-85473 virus were reached by 4 dpi (**Figure 1C**). According to previous studies with the viral C-85473 background [29, 30], fluorescence microscopy showed that ΔSH-C-85473 and ΔG-C-85473 viruses harbored high viral spread and hyperfusogenic phenotypes, comparable to the WT rC-85473 (**Figure 1D**).

Considering the different phenotypes of rHMPV virus derived from the A1/C-85473 and B2/CAN98-75 strains and the “attachment” role attributed to the G protein, we further evaluated the specific impact of the G and SH gene deletions on the binding and the entry of the rHMPV to LLC-MK2 cells (**Figure 2**). While the rCAN98-75 virus showed progressive binding kinetics, reaching a maximum of 68% GFP-positive cells after 3h of adsorption, the ΔSH-CAN98-75 virus showed faster binding in the first hour, yet achieving a maximal plateau of 47% by 2h (**Figure 2A**). Not surprisingly, the deletion of the G protein significantly hampered the binding capacity of the ΔG-CAN98-75 virus, with a mean of∼10% GFP-positive cells all throughout the experiment (**Figure 2A**). In contrast, and as observed for the replicative capacities, the binding ability of the C-85473-derived viruses was not significantly hampered at 3h by the gene deletions: 57% for ΔSH- and 54% for ΔG-C-85473 *versus* 43% for the WT rC-85473 (**Figure 2B**).

**Figure 2:**
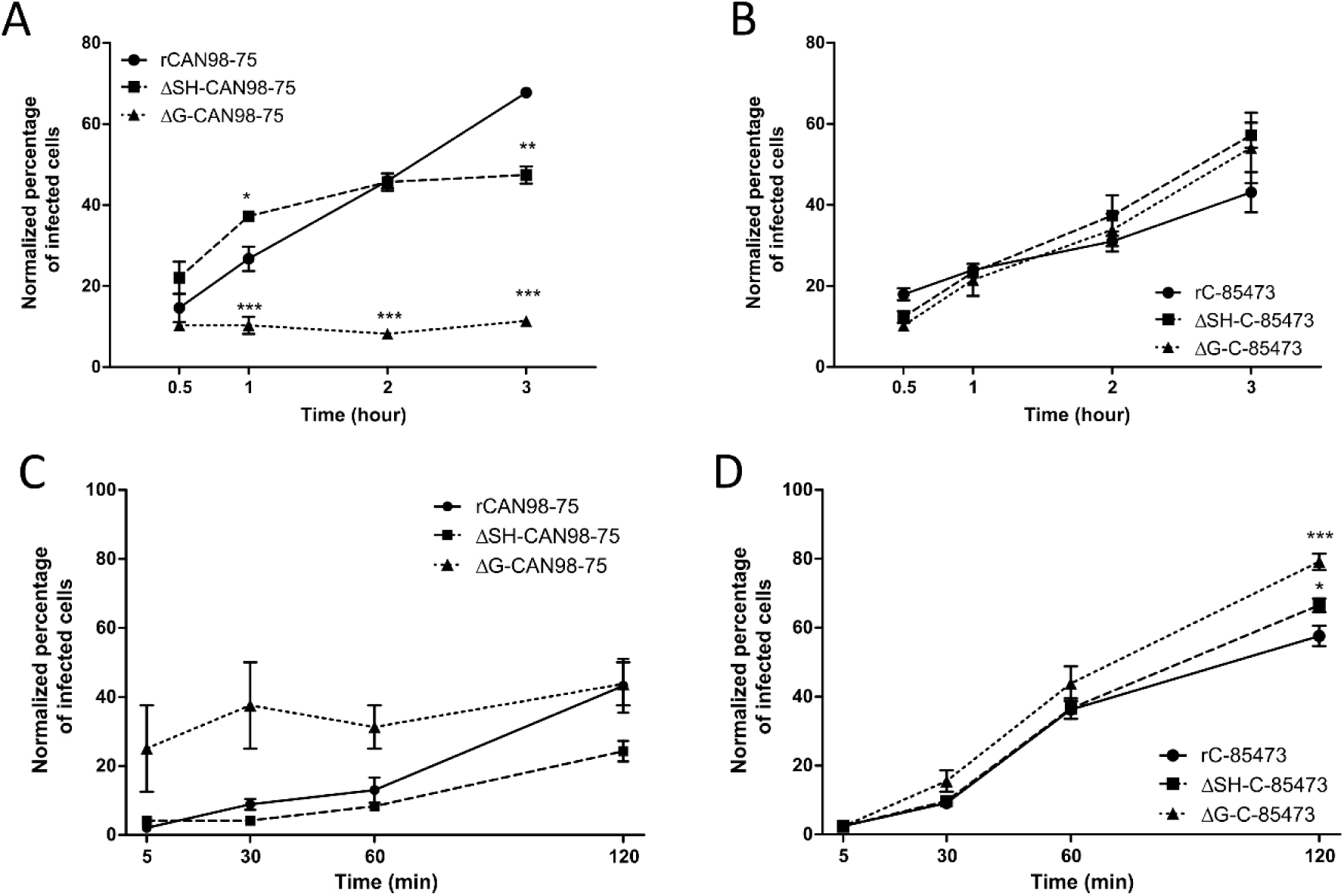
Binding and entry kinetics of WT, ΔSH- or ΔG-rHMPV viruses in LLC-MK2 cells. Binding kinetics of **(A)** rCAN98-75-derived viruses and **(B)** rC-85473-derived viruses or entry kinetics of **(C)** rCAN98-75-derived viruses and **(D)** rC-85473-derived viruses in LLC-MK2 cells were measured by flow cytometry as described in Materials and Methods. Mean values represented triplicates in 2 independent experiments. * p < 0.05, ** p < 0.01, *** p < 0.001 when comparing each ΔSH / ΔG virus to its WT counterpart using Repeated Measures Two-way ANOVA.

Regarding cell entry kinetics, the ΔSH-CAN98-75 virus seemed to enter LLC-MK2 cells slower than its WT counterpart and the ΔG-CAN98-75 virus, with 24% of ΔSH-CAN98-75-infected cells after 120 min compared to 43% for WT and ΔG-CAN98-75 viruses (**Figure 2C**). Conversely, the three rC-85473-derived viruses showed rather rapid cell entry kinetics with more than 36% of infected cells after 60 min. Interestingly, both ΔSH- and ΔG-C-85473 viruses showed higher entry properties compared to their WT counterpart after 120 min with 66%, 79% and 57% of infected cells, respectively (**Figure 2D**).

Altogether, our *in vitro* results on virus replication, cell binding and entry kinetics, suggest that G and SH deletions have strong differential impact on HMPV phenotypes depending on the viral background, hence highlighting a strain-dependent role of the viral surface proteins.

### ΔG- and ΔSH-viruses harbor different replicative properties in reconstituted human airway epithelium

We further investigated the properties of ΔG- and ΔSH-viruses by using reconstituted human airway epithelium (HAE) as a more physiological model of respiratory infection. Indeed, we previously showed that this model is permissive to HMPV infection and globally mimics the *in vivo* host respiratory epithelium response to such infection [36]. In line with these results, we observed that both rCAN98-75 and rC-85473-derived viruses are also able to infect, replicate and spread within the HAE (**Figure 3**).

**Figure 3:**
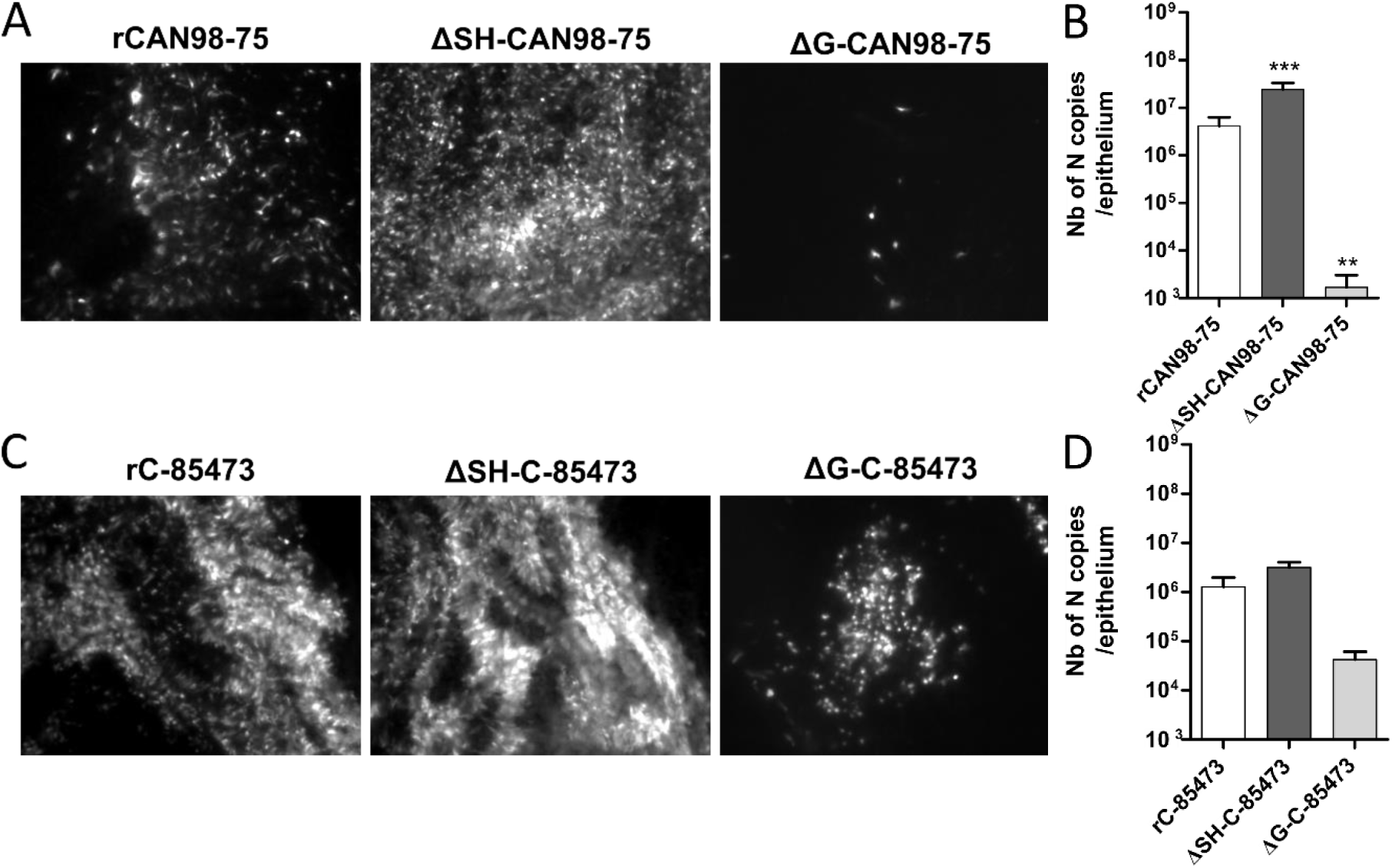
Recombinant HMPV viruses differ in infectivity and viral growth in reconstituted human airway epithelium (HAE). MucilAir™ epithelium from healthy donors were infected with the different rHMPV strains at a MOI of 0.1. At 5 dpi, the viral spread in infected ciliated cells was monitored by fluorescence microscopy (images at 10x magnification) and the quantity of viral genome was measured by specific RT-qPCR against the N viral gene from epithelium lysates, respectively (**A**) and (**B**) for rCAN98-75-derived viruses or (**C**) and (**D**) for rC-85473-derived viruses. Genome quantifications are shown as means ± SD and represent experimental triplicates. ** p < 0.01, *** p < 0.001 when comparing each ΔSH / ΔG virus to its WT counterpart using Repeated Measures Two-way ANOVA.

Based on the GFP expression pattern within the infected HAE at 5 dpi, we observed that the ΔSH-CAN98-75 spread more efficiently than its WT counterpart, at the difference of the ΔG-CAN98-75 virus, which appeared dramatically impaired (**Figure 3A**). These results were confirmed by the quantification of N gene copies number within the infected epithelium. In line with the fluorescent microscopy observations, the viral quantifications at 5 dpi indicated a significant 6-fold higher viral replication of the ΔSH-CAN98-75 virus and 3.3 log10 lower values for the ΔG-CAN98-75 virus in comparison to the WT CAN98-75 virus (**Figure 3B**). At the difference, the ΔSH-C-85473 virus acted similarly to its WT counterpart, considering both spread pattern and viral replication within the HAE model (**Figure 3C-D**). Besides, the spread of the ΔG-C-85473 virus seemed to be more affected in HAE model than in LLC-MK2 monolayer (**Figure 3C**) even though its replication diminished slightly in comparison to the WT rC-85473 virus, considering the 30-fold decrease of N gene copies number (**Figure 3D**).

These results indicate that rHMPV harbour different strain-dependent replicative properties in HAE model (**Figure 3**) compared to the LLC-MK2 model (**Figure 1**). In the CAN98-75 viral background, the SH deletion significantly enhances the viral replication and spread in HAE, whereas it causes a deleterious delay of replication in LLC-MK2 cells. In addition, the G deletion appears to attenuate excessively the viral replication in HAE, in line with results in LLC-MK2 model. In contrast, rC-85473 viruses show similar characteristics in both HAE and LLC-MK2 models, including comparable replication phenotypes between the WT virus and its derived deleted viruses, and especially the ΔSH-C-85473 virus.

Altogether, considering the characteristics of the rC-85473-derived viruses in LLC-MK2 and HAE models, we focused on this viral backbone to further investigate *in vivo* functional impact of SH- and G-deletions and evaluate their potential in the development of LAV candidates.

### Immunization with deleted-C-85473 viruses reduces HMPV disease severity in challenged BALB/c mice

We therefore infected mice intra-nasally with 5×10^5^ TCID_50_ of either ΔG-C-85473 or ΔSH-C-85473 viruses, the equivalent non-lethal dose inoculum needed to induce a significant (>10%) weight loss with the WT rC-85473 virus at 7 dpi (**Figure 4A**). Similar to the mock (non-infected) group and in contrast with the WT rC-85473, neither weight loss nor clinical signs were observed in the ΔG-C-85473 and ΔSH-C-85473-infected groups during the 14-day follow-up. In contrast, lung viral titers at 5 dpi were comparable between WT and deleted viruses, as determined by both cell culture and molecular methods (**Figures 4B-C**). In agreement with the weight curves, lower histopathological scores were recorded for both ΔG-C-85473 and ΔSH-C-85473-infected mice compared to the WT group (1.8 and 2.4 versus 5.3, respectively), particularly owing to significant reductions in interstitial, perivascular and/or intra-alveolar inflammation (**Figure 4D**).

**Figure 4:**
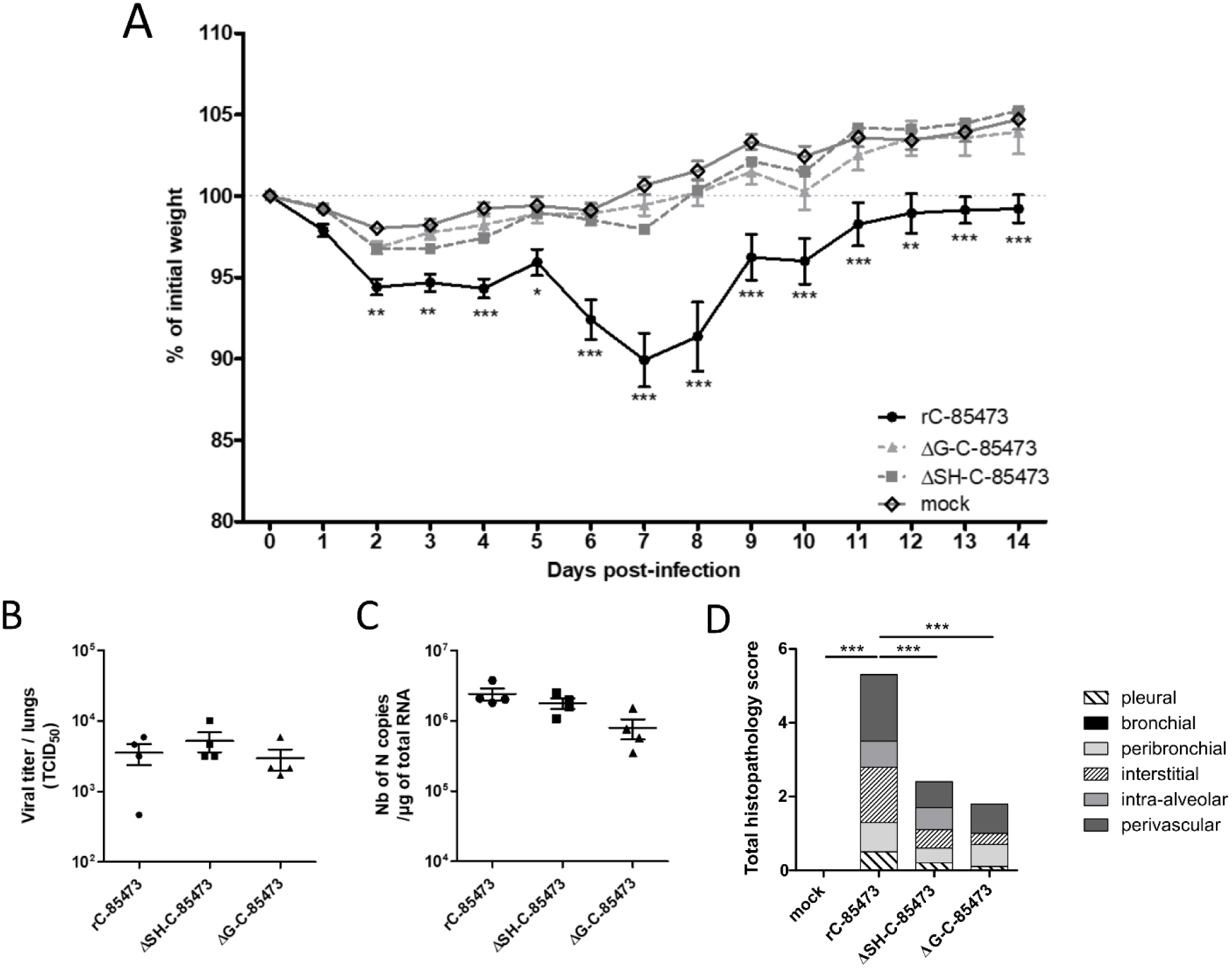
Weight loss, lung viral titers and histopathological scores of BALB/c mice infected with rC-85473-derived LAV candidates. BALB/c mice were intranasally infected with 5×10^5^ TCID_50_ of rC-85473 rHMPV viruses (WT, ΔSH and ΔG). (**A**) Weight loss was monitored during 14 days since infection (n = 16). At day 5 pi, mice were euthanized and lungs were harvested to measure viral titers by both TCID50 titration (**B**) and quantitative RT-PCR (**C**) (n = 4). (**D**) Cumulative histopathological scores (bronchial, peribronchial, perivascular, interstitial, pleural and intra-alveolar inflammation scores) of other infected mouse lungs were evaluated on day 5 pi (n=5). Data are shown as means ± SEM. *, p < 0.05, **, p < 0.01, ***, p < 0.001 when comparing each ΔSH / ΔG virus to its WT counterpart using Repeated Measures Two-way ANOVA.

We then evaluated the capacity of ΔSH- and ΔG-C-85473 attenuated viruses to protect mice from a lethal viral challenge. Upon viral challenge with 1×10^6^ TCID_50_ of rC-85473 virus, mock-immunized mice showed significant (50%) HMPV-associated mortality, starting on day 5 post-challenge and spanning until day 8 post-challenge, with mean weight loss of 11% (peak after 7 days) (**Figure 5A**). Conversely, both ΔSH- and ΔG-C-85473-immunized mice groups, as well as control rC-85473-immunized mice, were completely protected from HMPV-associated mortality induced by viral challenge and showed mean maximum weight losses of 3-4% of their initial weight (peak at day 2 post-challenge) (**Figure 5A-B**).

**Figure 5:**
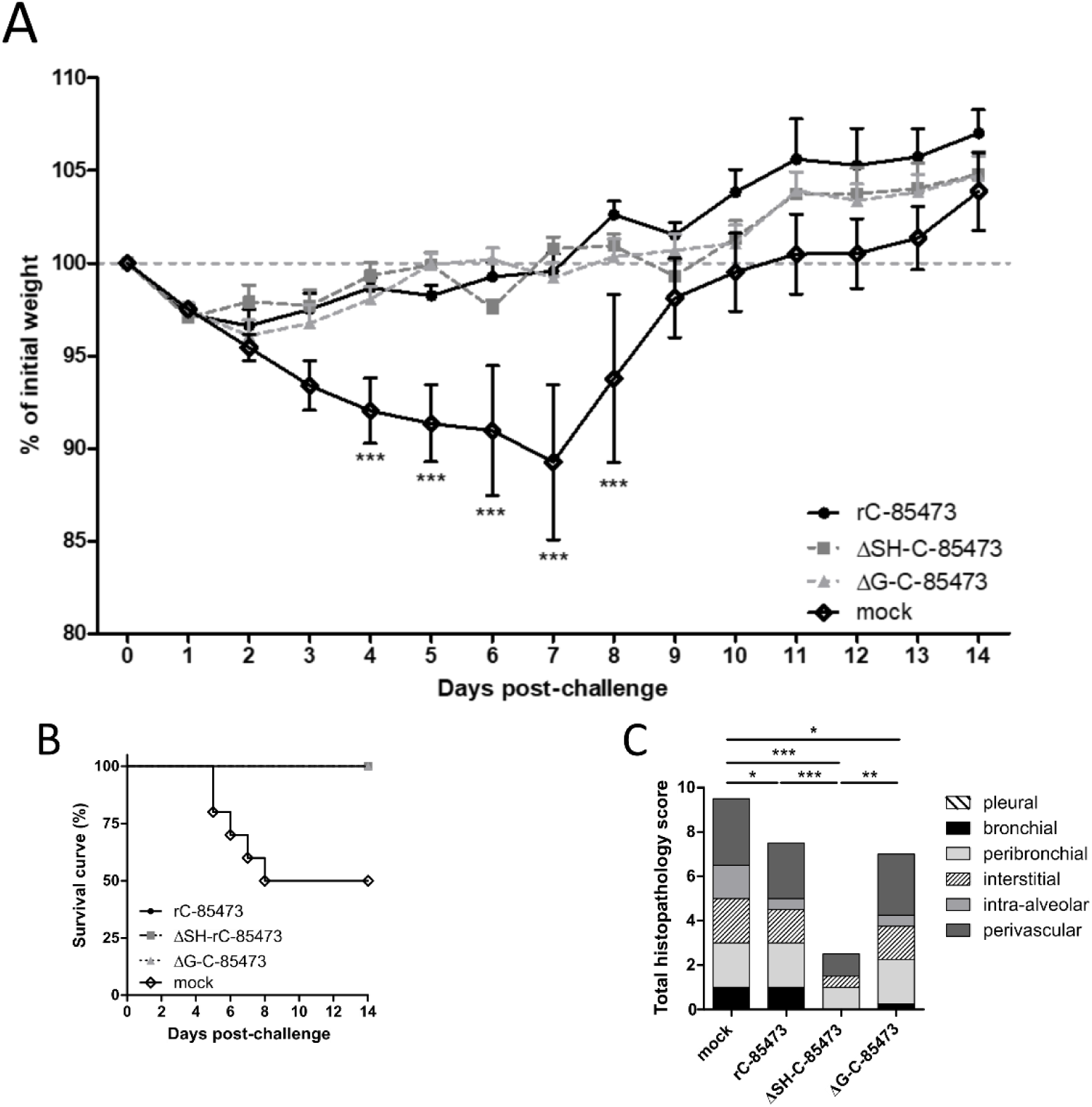
Weight loss, survival and histopathological scores of HMPV-immunized mice after viral challenge. BALB/c mice were intranasally immunized with 5×10^5^ TCID_50_ of ΔSH-C-85473, ΔG-C-85473 or WT rC85473 viruses (immunization control), as previously described. Three weeks after immunization, animals (n=16) were inoculated with 1×10^6^ TCID_50_ (LD_50_) of rC-85473 (viral challenge). **(A)** Weight loss and (**B**) mortality were monitored during 14 days after the viral challenge (n=10). (**C**) Cumulative histopathological scores (bronchial, peribronchial, perivascular, interstitial, pleural and intra-alveolar inflammation scores) of infected mouse lungs were evaluated on day 5 post challenge (n=2) while pulmonary viral titers were measured by RT-qPCR (n = 4, **Table 2**). Data are shown as means ± SEM. * p < 0.05, ** p < 0.01, *** p < 0.001 when comparing each ΔSH / ΔG virus to its WT counterpart using Repeated Measures Two-way ANOVA.

Similarly to the rC-85473 WT virus, the protection induced by immunization with ΔG-C-85473 and ΔSH-C-85473 was associated with the induction of high neutralizing antibody titers at 21 days after challenge, particularly in the case of ΔSH-C-85473 (**Table 2**). Importantly, neutralization assays also showed that induced antibody responses were effective in neutralizing WT patient-derived strains as well as the heterologous CAN98-75 HMPV B strain (**Table 2).** In line with these results, we were unable to recover HMPV viruses or detect viral genome from the lungs of any of the three different rHMPV-immunized groups on day 5 post-challenge (**Table 2**). However, we still recorded reduced but significant pulmonary inflammation in WT- and ΔG-C-85473-immunized mice, compared to mock-immunized mice (scores of 7.5 and 7 versus 9.5, respectively), whereas ΔSH-C-85473-immunized mice had a much lower histopathological score of 2.5 (**Figure 5C**). Indeed, ΔSH-C-85473-immunized group showed reduced interstitial and perivascular inflammations and no bronchial and intra-alveolar inflammation (**Figure 5C**). In contrast to mock-immunized mice, none of the rHMPV-immunized groups developed pulmonary edema after viral challenge (**Figure A1**).

**Table 2.** Immunogenicity and protective efficiency of HMPV C-85473-derived viruses in mice.

| Imm. virus | Imm.<br>Inoculum<br>(log10<br>TCID<sub>50</sub>) | Reciprocal neutralization titer 21 days post-challenge against different HMPV strains (n=4) <sup>1</sup> |  |  | Challenge virus replication in lungs<br>5 days post-challenge<br>(n = 4) |  |
| --- | --- | --- | --- | --- | --- | --- |
|  |  | rC-85473 | WT C-85473 | WT CAN98-75 | Mice with<br>detectable<br>virus (%) | Mean virus<br>titer (log10 N<br>copies/μg<br>RNA) |
| mock |  | 10 | 7.5 | 12.5 | 100 | 4.8 ± 4.3 |
| rC-85473 | 5.7 | > 160 | 80 | 60 | 0 | < 1 |
| ΔSH-C-85473 | 5.7 | > 160 | > 160 | > 160 | 0 | < 1 |
| ΔG-C-85473 | 5.7 | > 160 | 80 | 70 | 0 | < 1 |

<sup>1</sup> Sera were tested for neutralization efficiency against rC-85473 virus, which is the same recombinant virus used for infectious challenge; WT C-85473, as homologous A1 virus; and WT CAN98-75, as heterologous B2 strain. Two biological replicates per group were tested. The “naïve” status of mice was confirmed by neutralizing antibody dosage of sera harvested one day before intranasal immunization (reciprocal titer <5). Imm. = immunization.

To further characterize the immune response after viral challenge of the rHMPV-immunized mice, we characterized the production of pulmonary cytokines/chemokines and the subsequent infiltration of different immune cell populations into the lungs. As soon as 1 day after the challenge, we measured high expression levels of cytokines, corresponding to an acute response to high dose of viral challenge, as it is the case for RANTES (19 000pg/ml, **Figure 6A**). Interestingly, higher levels of several cytokines/chemokines (IL-10, IL-6, G-CSF, TNF-α) were measured in the lungs of ΔSH-immunized mice in comparison with WT and ΔG-immunized groups (**Figure 6A**). In accordance with the absence of detectable virus (**Table 2**) and the decreased histopathological scores (**Figure 5C**), the level of most of the measured cytokines lowered 5 days post-challenge (**Figure 6A**). In addition, we observed an important recruitment of leukocyte populations in rHMPV-immunized groups from day 1 after challenge, which persisted for at least five days later (**Figure 6B**). Of note, the ΔSH-immunized group showed significant enhanced infiltration of leukocytes in lungs 5 days post-challenge, particularly T CD4+, T CD8+, and B cell populations (**Figure 6B**).

**Figure 6:**
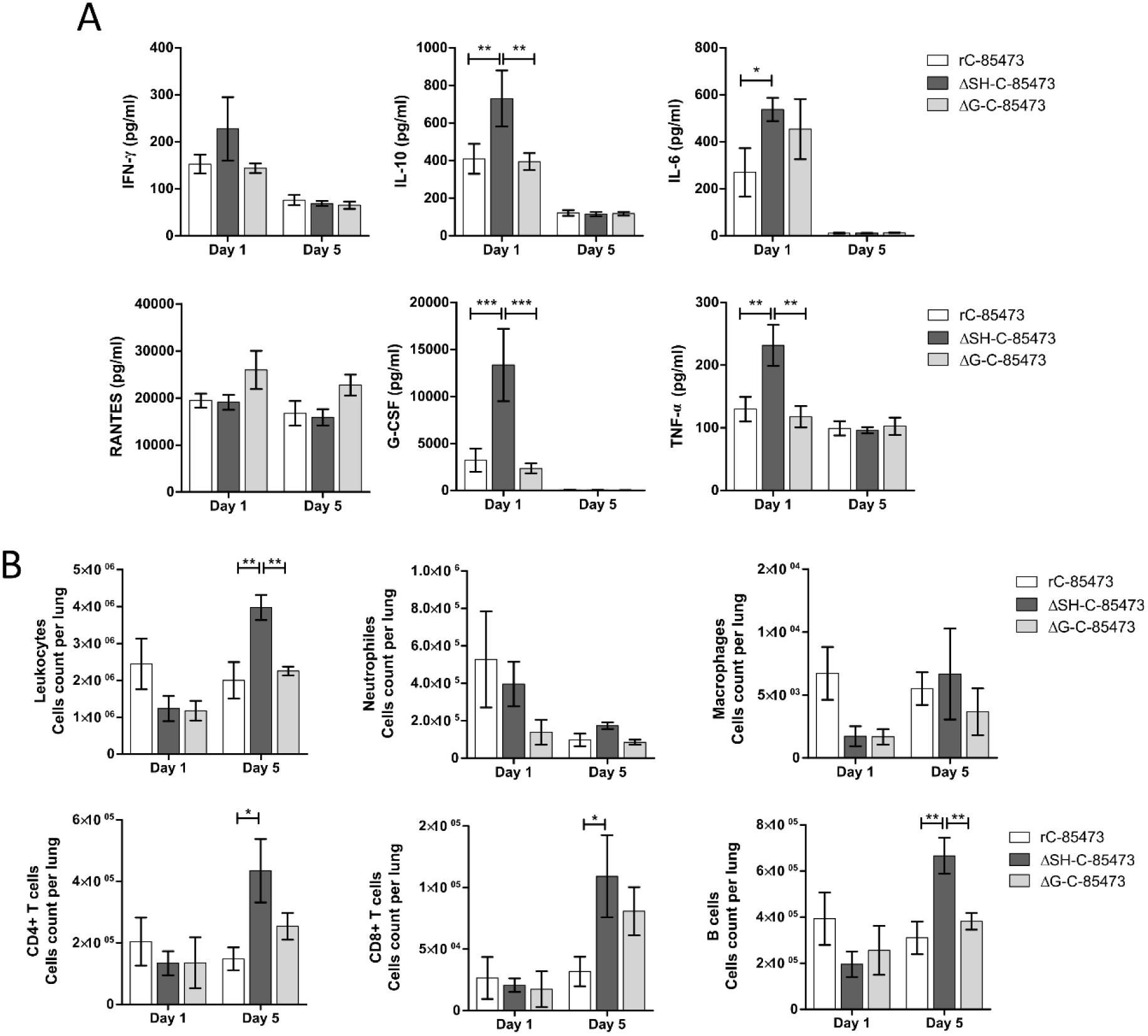
HMPV-immunized mice showed distinct cytokine/chemokine and immune cell infiltration profiles through time after viral challenge. BALB/c mice were intranasally immunized with 5×10^5^ TCID_50_ of ΔSH-C-85473, ΔG-C-85473 or WT rC85473 viruses (immunization control), as previously described. Three weeks after immunization, animals were inoculated with 1×10^6^ TCID_50_ (LD_50_) of rC-85473 (viral challenge). **(A)** Pulmonary inflammatory cytokine/chemokine levels in HMPV-immunized challenged mice were determined from lung homogenates on days 1 and 5 post-challenge, using multiplex bead-based assay. Six cytokines/chemokines (IFN-γ, IL-10, IL-6, RANTES, G-CSF, TNF-α) were selected considering both their detection levels and significance in HMPV-related disease. **(B)** Immune cell infiltration in lung homogenates was evaluated by flow cytometry on days 1 and 5 post challenge. * p < 0.05; ** p < 0.01; *** p < 0.001 comparing each group using Repeated Measures Two-way ANOVA.

Overall, our results suggest that deleted rC-85473 viruses harbor distinct but attenuated and protective properties in the BALB/c model against a lethal HMPV viral challenge, together with reduced disease severity, weaker inflammatory responses and a balanced stimulation of the immune response, especially in the case of the ΔSH-C-85473 virus.

## 4. Discussion

Despite the necessity to develop vaccination strategies to prevent ARTI, neither FDA-approved vaccine nor specific antiviral against HMPV are currently available [37, 38]. Considering the major burden of this virus in infants and young children, HMPV live-attenuated vaccines offer interesting properties to protect this specific population [8]. In that regard, while the first and only phase I clinical trial based on human-to-avian HMPV P substitution was non-conclusive [39], attenuation approaches based on gene deletion showed promising results in animals [27, 40]. Nevertheless, previous studies on HMPV LAV were mainly restricted to the A2/CAN97-83 strain and did not take into account the genetic variability of HMPV viruses between and within subgroups, especially regarding the poorly conserved G and SH glycoproteins [41, 42]. In fact, the limited knowledge on HMPV virulence factors and their putative roles still hampers the development of efficient therapeutic and prophylactic strategies against severe HMPV disease. In this study, we sought to address the problematic of the impact of viral background on the differential effect of G and SH deletions by using two patient-derived HMPV strains, A1/C-85473 and B2/CAN98-75. We demonstrated that G- or SH-deleted A1/C-85473 viruses had unchanged replicative capacity *in vitro* compared to their WT counterpart (**Figure 1**), in line with a previous study advocating the putative “accessory role” of these two gene-products [27]. However, we observed that viral replication in LLC-MK2 cells was significantly decreased when G and, to a lesser extent SH, were deleted from the B2/CAN98-75 virus (**Figure 1**). Such a distinct replication phenotype between A and B viruses (A1/C-85473 and A2/CAN97-83 *vs* B2/CAN98-75) suggests that the HMPV genetic background might influence the impact of gene deletions on viral fitness. This is of particular importance if we consider efficient production yield (i.e. comparable to that of the WT virus) as a major requirement for the development of a LAV candidate.

We also previously showed that C-85473 and CAN98-75 strains induce very different fusogenic phenotypes *in vitro* and pathogenicity *in vivo*, thanks to the specific characteristics of their respective F glycoproteins [29, 30]. Besides its major role in viral fusion, the F protein may also retain importance for viral binding and entry into cell [17, 20, 21]. In this way, the C-85473 virus appeared to be less dependent on the G protein to perform virus-to-cell binding than the CAN98-75 virus (**Figure 2**). It seems therefore possible that the lack of the G protein could be compensated by the hyperfusogenic activity of the C-85473 F protein. In our study, the differential profiles exhibited by the two strains strengthen this hypothesis, as illustrated by the enhanced entry kinetics of the rC-85473 virus in comparison to the rCAN98-75 virus, despite the improved capacity of the latter to bind LLC-MK2 cells (**Figure 2**). Alternatively, although its specific function is still debated, SH is presumed to be a putative cofactor of some F proteins and therefore it could participate in the viral entry [26]. In this way, SH deletion could lead to mild viral attenuation *in vitro*, as reported for the ΔSH-CAN98-75 virus in **Figure 1**.

Contrasting with the LLC-MK2 model, the replication and spread of both ΔG viruses was more limited in the HAE model, whereas the ΔSH viruses showed efficient infectivity, notably in the case of the ΔSH-CAN98-75 virus (**Figure 3**). Such divergent phenotypes between deleted viruses was previously shown with HRSV, for which the G protein differently contributes to virus-cell binding depending on both the cell lines and the virus strains studied [43]. In light of our results, we therefore propose that the stratified nature and surface receptor configuration of the physiological HAE model, combined with the presence of cilia and mucus, lead to a more limiting environment for effective infection with a G-deleted HMPV virus. Taken together, our data show for the first time the importance of HMPV background and experimental models which must be considered in the study of G and SH protein functions.

Based on their specific characteristics in the LLC-MK2 and HAE models, we focused on the two deleted (ΔSH and ΔG) viruses issued from the C-85473 strain for further investigation in BALB/c mice. At the difference of other small animal models previously used, the murine model allows the monitoring of weight loss, lung viral replication/inflammation and other clinical signs during the time course of HMPV infection [44]. In our study, the ΔSH-C-85473 and ΔG-C-85473 viruses replicated in mice lungs similarly to their WT counterpart at 5 dpi while inducing no weight loss or mortality and lesser tissue inflammation (**Figure 4**). Accordingly, we recently showed that the SH deletion in the C-85473 backbone limits the virus-induced activation of NLRP3-inflammasome both *in vitro* and *in vivo*, and subsequently reduces inflammation and pathogenicity in HMPV infected mice [34].

Moreover, ΔSH-C-85473 and ΔG-C-85473 viruses induced the efficient production of HMPV-specific neutralizing antibodies; fully protecting mice against viral challenge with rC-85473 (**Figure 5, Table 2**). Remarkably, this antibody response was also effective against a heterologous HMPV B strain (**Table 2)**. Additionally, ΔSH-C-85473-immunized mice showed a distinct pulmonary cytokine/chemokine profile after viral challenge, with a significant increasing expression of several HMPV-response related markers compared to the WT and ΔG-C-85473 immunized groups (**Figure 6A**). Interestingly, a high expression of IL-10 cytokine [45] and G-CSF chemokine, which were reported as T-cell activators [46], was concomitant with an enhanced recruitment of immune cells (T CD4+, T CD8+, B cells) in lungs of ΔSH-C-85473-immunized mice at 5 days post-challenge (**Figure 6B**). Moreover, such a production of anti-inflammatory cytokine/chemokine counteracts the observed increased production of TNF-α and IL-6 pro-inflammatory (**Figure 6A**).

In conclusion, given the significant clinical burden of HMPV [1, 2] and some pitfalls in the development of vaccine candidates [7, 39, 47], we considered important to reexamine the gene deletion-based attenuation approach [6, 27, 40]. Our study provides compelling evidence on the so far overlooked variable function of HMPV G and SH proteins relative to the viral background, the differential effects resulting from their deletion and the potential implications for the design of LAV candidates. In line with this rationale, the attenuated and broad protective properties displayed by the ΔSH-C-85473-engineered virus, yet not inducing enhanced disease markers, in addition to its efficient replication in cell-based system, make it a promising live-attenuated HMPV vaccine candidate.

## 5. Patents

The authors declare a patent submission (FR1856801, PCT FR2019/051759) about HMPV LAV candidates.

## Supplementary Materials

Figure A1: HMPV-immunized mice showed significant differences in histopathology 5 days after viral challenge.

## Author Contributions

Conceptualization, J.D., M.R-C. and G.B.; methodology, J.D. and M-H.C.; validation, J.D. and M-E.H.; formal analysis, J.D and C.NdL.; investigation, J.D., B.P., O.U., M-C.V., J.C., A.T., T.J., A.P. and C.C.; resources, J.C. and A.T.; writing—original draft preparation, J.D.; writing—review and editing, A.P., O.T., M-E.H., M.R-C. and G.B.; visualization, J.D. and C.NdL.; supervision, B.L., G.B. and M.R-C.; project administration, J.D. G.B. and M.R-C; funding acquisition, G.B. and M.R-C.

## Acknowledgments

We thank the microscopy and flow cytometry services of the Plateforme of Bio-Imagerie of CRI in Québec and the Centre d’Imagerie Quantitative Lyon-Est (CIQLE) in Lyon, as well as the animal care services of the Plateau de Biologie Expérimentale de la Souris in Lyon.

