## Supplementary figure 1 for "Strain-dependent impact of G and SH deletions provide new insights for live-attenuated HMPV vaccine development"

**Figure A1. HMPV-immunized mice showed significant differences in histopathology 5 days after viral challenge.**

After intranasal immunization with $5x10^5$ $TCID_{50}$ of ΔSH-C-85473, ΔG-C-85473 or WT rC85473 virus, mice were inoculated with $1\times10^6$ $TCID_{50}$ ($LD_{50}$) of rC-85473 3-weeks later. Tissues observations (x100 magnification) and measure of pulmonary edema scores were performed on mice lungs on day 5 post-challenge (n=2).

| | | pulmonary edema score |
|---|---|---|
| challenged mice mock-immunized | 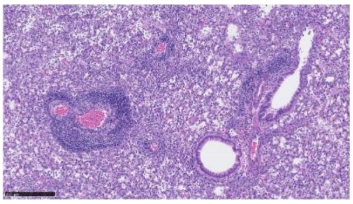 | 1.5 |
| challenged mice rC-85473 WT immunized | 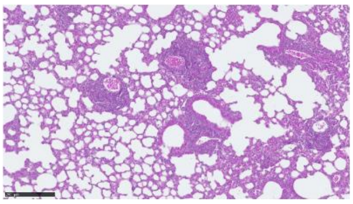 | 0 |
| challenged mice ΔSH-C-85473 immunized | 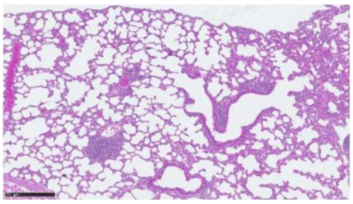 | 0 |
| challenged mice ΔG-C-85473 immunized | 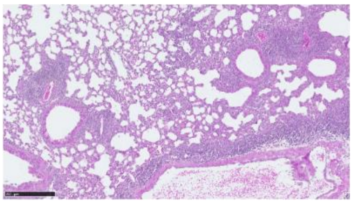 | 0 |
